# Emotional responsiveness task in emotional distress: correlated of functional neuroimaging in anorexia and bulimia

**DOI:** 10.1101/2020.03.31.018424

**Authors:** Federico D’Agata, Paola Caroppo, Angela Spalatro, Luca Lavagnino, Giovanni Abbate Daga, Andrea Boghi, Mauro Bergui, Alessandro Cicerale, Benedetto Vitiello, Secondo Fassino, Birgit Derntl, Federico Amianto

## Abstract

**Aim:** The present study aims to extend the knowledge of the neural correlates of emotion processing in first episode subjects affected by anorexia nervosa (AN) or bulimia nervosa (BN). We applied an emotional distress paradigm targeting negative emotions thought to be relevant for interpersonal difficulties and therapeutic resistance mechanisms.

**Methods:** The current study applied a neuroimaging paradigm eliciting affective responses to 44 female participants with newly diagnosed AN or BN and 20 matched controls. The measurements also included an extensive assessment comprised of clinical scales, neuropsychological tests, measures of emotion processing and empathy.

**Results:** AN and BN did not differ from controls in terms of emotional response, emotion matching, self-reported empathy and cognitive performance. However, scores of eating disorder and psychopathological clinical scores, as well alexithymia levels, were increased in AN and BN. On a neural level, no significant group differences emerged, even when focusing on a region of interest selected a priori: the amygdala.

**Conclusions:** Our data are against the hypothesis that participants with AN or BN display a reduced emotional responsiveness. This supports the hypothesis that relational difficulties, as well as therapeutic resistance, are not secondary to simple difficulty in feeling and identifying basic negative emotions in AN and BN participants.

## 1. Introduction

Anorexia nervosa (AN) and Bulimia nervosa (BN) are the two major Eating Disorders (ED): serious and complex psychiatric conditions with a multifactorial biopsychosocial pathogenesis often characterized by a chronic and disabling course and only partial therapeutic success [1,2]. Young girls are especially affected by AN, which is the pathology with the highest mortality risk and the lowest response to treatment across ED [3]. Prioritizing the treatment of symptoms results in better outcomes in BN and allows dealing with the main cause of mortality in AN [4]. It remains controversial whether doing so ignores core psychopathological elements, linked to more complex symptoms and long-term outcomes such as relationship difficulties or impairments in affect regulation, reflective functioning, and coherence of mind [5]. Psychotherapeutic treatment often focuses on these aspects and therapists are frequently faced with marked difficulty in engaging subjects affected by AN and maintaining treatment adherence [6]. In BN the difficulties are related to coping with high emotional arousal when facing social and affective stimuli. These difficulties also challenge a complex therapeutic approach [7].

The problems of individuals affected by ED in social interactions and psychotherapeutic engagement are indicators of serious difficulties in the management of interpersonal relationships, as well as emotional dysregulation [8]. Although not included in the current diagnostic criteria of ED (DSM-5 and ICD-10), emerging evidence points to deficits in socioemotional functioning [9]. Consequently, several modern therapeutic models incorporate the role of emotional difficulties, social anxiety and poor social support in the maintenance of the disorder [8].

Empathy represents a core function for social coherence and building relationships [10]. Based on the abovementioned socio-emotional difficulties and related problems in ED, one may assume that empathy is systematically altered in ED and its impairment potentially represents a relevant risk factor. Several studies applied self-reported empathy measures or assessed emotion recognition performance but reported mixed results [11–14]. Thus, it is unclear how much emotion processing, empathy and social competencies are affected in ED and what mechanisms mediate the insurgence of relational difficulties.

The majority of neuroimaging studies in the field of ED have investigated the neurobiological correlates of body shape, reward and food stimuli [15,16], while the number of studies focusing on emotion/empathy is still scarce and almost limited to the functional magnetic imaging (fMRI) correlates of implicit and/or explicit face emotion processing in patients affected by AN [17–22]. In BN much less is known regarding the neural circuits underlying emotion processing [16] and hardly any neuroimaging study adopted a transdiagnostic approach by including both AN and BN.

The present study aimed to extend the knowledge of the neural correlates of emotion processing in first-episode young women affected by AN or BN, using an emotional response fMRI paradigm focusing on negative emotions, as these could be relevant for interpersonal problems and therapeutic resistance mechanisms [8]. The importance of selecting first-episode participants derives from the aim of excluding secondary effects of the disorders and of prolonged or repeated therapeutic interventions on brain functioning. We compared performance of women affected by AN or BN and a group of matched healthy controls with an extensive assessment that included clinical scales, neuropsychological tests, and self-report questionnaires of emotion processing and empathy and with fMRI. Based on previous studies we hypothesized: i) behavioral and/or self-reported differences in emotion processing/empathy measures between the three groups, ii) group differences for the activation of specific limbic areas known to be highly involved in emotional processing and empathy, i.e. amygdala [23–26], and iii) a critical contribution of neuroimaging data to distinguish between the three groups as compared to the behavioral or self-report measures, as it should be a less biased and more sensitive measurement tool.

## 2. Materials and Methods

### 2.1 Sample

Twenty-five female individuals affected by AN (20 restricting, 5 binge/purging) and 19 by BN (15 purging, 4 not purging), were enrolled from the outpatient service of the Pilot Centre for the Diagnosis and Treatment of Eating Disorders of the Department of Neuroscience, “AOU Città della Salute e della Scienza” of Turin, Italy. ED was diagnosed using the Structured Clinical Interview for DSM-IV-TR (SCID). The inclusion criteria for the study were: female sex; age 16-30 years; right-handedness (assessed by Edinburgh Handedness Inventory); body mass index (BMI) from 15.0 to 17.5 for AN and from 19.0 to 25.0 for BN; no past or present mental disorder except for the current ED first-episode; no axis II disorders (assessed by SCID II, [27]; no current or past pharmacological treatment; no drug or alcohol abuse; no history of diabetes or other somatic diseases, no past or present psychotherapy treatment and duration of symptoms shorter than 2 years. From a global sample of 109 assessed ED participants, only 44 met the inclusion criteria and were finally enrolled in the present study.

Twenty healthy women were recruited as controls (CN) through local advertisement. The inclusion criteria for the CN group were like those for BN (BMI range 19.0-25.0), except for having neither a current nor lifetime mental disorder.

All participants gave their written informed consent to the study. For minors (3 AN, 1 BN), written informed consent was obtained from parents. The study was approved by the local Ethics Committee [Comitato Etico Interaziendale A.O.U. Città della Salute e della Scienza di Torino - A.O. Ordine Mauriziano - A.S.L. Città di Torino, approval #12042010].

### 2.2 Clinical and self-report data

The clinical assessment included the *Eating Disorder Inventory* 2 (EDI-2) [28], the *Symptom Checklist-90* (SCL-90) [29], the *Empathy Quotient* [30] and the *Toronto Alexithymia Scale* (TAS-20) [31]. Detailed information on the scales can be found in the supplementary material (section S1.1).

### 2.3 Neuropsychological tests

All participants performed a comprehensive neuropsychological testing battery assessing attention, memory and executive functions to account for possible interference of cognitive differences in emotional processing among groups. Detailed information can be found in the supplementary material (section S1.1).

### 2.4 MRI acquisition

MRI data were collected on a Philips Achieva 1.5T scanner. First, participants underwent the affective responsiveness task that comprised 480 continuous gradient-recalled EPI volumes (TR=2300ms, TE=40ms, FA=90°, 30 axial slices, matrix=128×128, slice thickness=4mm, no gap, voxel size=1.8×1.8× 4mm^3^, field of view=23cm). After fMRI, anatomical high-resolution images were acquired using T1-weighted 3D Turbo Field-Echo sequence (matrix=256×256, 190 contiguous sagittal slices, TR=7ms, TE=3ms, TFE shots=89, voxel size 1×1×1mm^3^).

#### 2.4.1 Affective responsiveness fMRI paradigm

We used a modified and shortened version of a previously tested fMRI paradigm [32–35]. Forty short written sentences were presented, describing real-life situations inducing anger, fear, disgust (e.g. ‘You are walking in a meadow and step on dog excrement’ for disgust) or containing a neutral content (e.g. ‘You are on the couch watching television’). As reported previously, stimuli were validated by independent female and male raters [32] and only stimuli that were clearly classified as belonging to one emotional category (>70%) were selected for the study. We presented 10 sentences per condition (disgust, anger, fear, neutral). Participants had to imagine how they would feel if they were experiencing those situations. We were particularly interested in the emotional response to distressful situations. Stimuli were presented for 7 sec. After emotion induction participants were presented with two facial expressions, one with the same emotion as the induction (correct) and the other with a match chosen randomly from the other options and asked to choose between the two. Participants had a maximum of 7 sec to respond. The response was followed by 7 sec of inter-trial-interval (cross fixation), then another trial started. The number of correct answers for the matching of emotional sentences and faces was recorded as a proxy of emotion processing (score 0-30). A right answer likely means a correct emotional response to the imaginary situation, correct identification of the emotions in the two displayed faces, and correct comparison and matching. After a neutral situation, participants were presented with two neutral faces, one male and one female, and they had to identify the female within 7 seconds. The number of correct answers was recorded as a proxy of the sustained attention and active participation to the task (score 0-10).

The order presentation of neutral and emotional stimuli was counterbalanced across participants. Figure 1 illustrates an example of the task. Further information on the experimental design can be found in the supplementary material (section S1.2).

**Figure 1:**
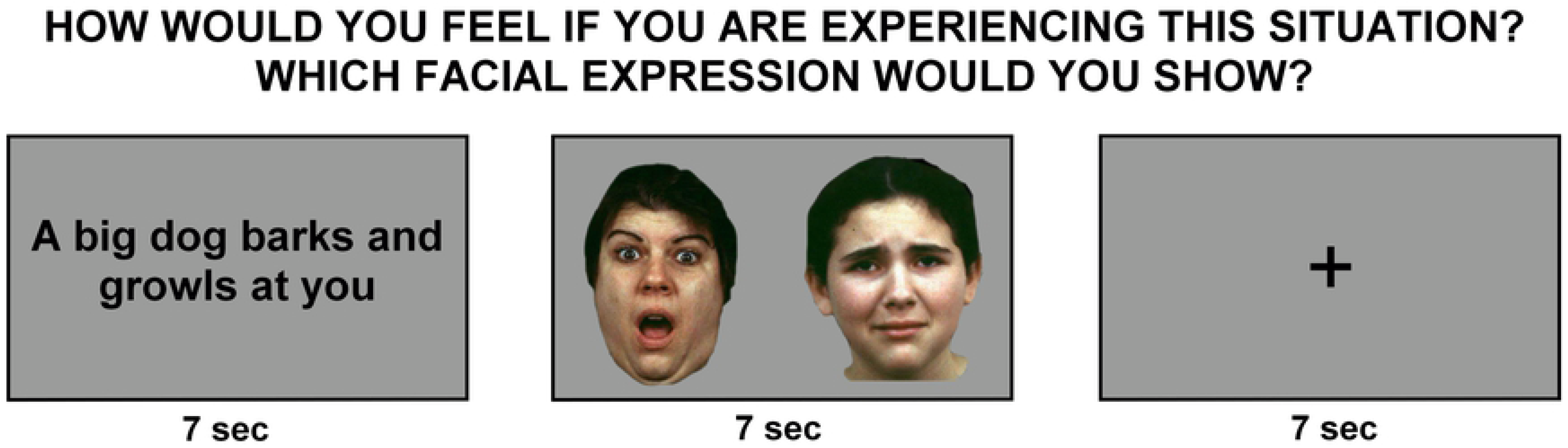
Illustration of the emotional response task. In every trial a short sentence with negative or neutral valence was presented, after which the participant was asked to choose between two stimuli (one correct and one wrong) and selected the facial expression that best matched the induced feelings. The paradigm consisted of 30 negative emotion trials (10 anger, 10 fear, 10 disgust) and 10 neutral trials.

### 2.5 Data analyses

#### 2.5.1 Statistical comparison of behavioral data

Statistical analyses were performed using IBM SPSS Statistics for Windows (Version 20.0. Armonk, NY: IBM Corp.) and level of significance was set to p<.05 corrected with False Discovery Rate (FDR q=.05) to account for multiple comparisons [36]. Behavioral and questionnaire data were compared using one-way ANOVAs with group as between-subjects factor. When needed, we performed a repeated measure ANOVA (rmANOVA) with emotion as within-subject factor. We used Tukey post-hoc tests to disentangle group differences when the main effect was significant. Effect sizes are reported using partial eta squared (η^2^).

#### 2.5.2 FMRI data processing

Functional data were preprocessed using SPM8 (http://www.fil.ion.ucl.ac.uk/spm) on MATLAB 7.5 environment. Images were motion corrected, normalized into the standardized stereotaxic MNI space using T1 coregistered images and spatially smoothed using an isotropic Gaussian kernel with a full-width-at-half-maximum (FWHM) of 8mm.

The estimation of head motion was tested with a procedure previously described by Yuan et al. [37]. The magnitude of head motion for six parameters (three for shift and three for rotation) was obtained for each participant and the averaged head motion parameters were calculated. An ANOVA was computed to test for differences between the groups. The result was non-significant, both for rotation (p=. 8; average movement × TR = 0.05º) and for shift (p=.6; average movement × TR=0.1mm). Yuan and colleagues suggest exclusion of participants with head motion exceeding 4 SDs (in our sample: rotation>0.1°; shift>0.2mm). None of our participants exceeded this threshold; therefore, data of all women were included in the analyses.

##### Task-based fMRI analyses

For this event-related design, each stimulus type was modeled with a separate regressor convolved with the canonical hemodynamic response function (HRF). Also, responses to the two different tasks were modeled as separate regressors. The participants gave correct answers in more than 85% of the trials, and the number of wrong responses did not differ significantly between groups (p=.5). The time-series movement estimated parameters (3 translations, 3 rotations) were included as covariates of no interest and the HRF time derivative was modelled.

Statistical analysis was performed with a two-step hierarchical estimation. At a first individual level, we saved three contrast images for every participant: disgust (D), anger (A), fear (F) compared to the neutral condition N (D>N, A>N, F>N, the contrasts compared 10 emotional trials with 10 neutral trials for each of the three conditions). We did not include in this model the response related regressors (button press), as we only focused on the neural activation during imagined situations. Contrast images from all participants were included in three second-level random-effects analysis to detect group differences, never mixing D, A and F or overrepresenting in the same model the N trials. We performed three one-way ANCOVA (General Linear Model) with GROUP as a between-subjects factor (AN, BN, CN) and controlling for atrophy. As cerebral atrophy could be a confounding factor in ED, as previously reported [38], we used VBM8 toolbox [39] to automatically extract gray matter (GM), white matter (WM) and cerebro-spinal fluid (CSF) volumes of all the participants. We compared parameters between groups including the significantly different brain volumes (CSF) as a covariate of no interest. Statistical inferences were performed by applying the Random Field Theory. Maps were thresholded at the p<.05 FWE cluster-level corrected (FWEc, meaning uncorrected p<.001, filtered for small clusters or cluster extent>FWE cluster size threshold).

##### Emotion ROI analyses

We performed region of interest (ROI) analyses on brain areas (left and right amygdala) that were selected a priori as they have been consistently reported as key regions in emotion processing. To do so, we used the MarsBaR region of interest toolbox for SPM (http://marsbar.sourceforge.net). ROIs mean signals were extracted using the AAL atlas delineated ROI in the MNI space. We used the same statistical ANCOVA models of the whole-brain analysis previously reported for every ROI separately. We applied Bonferroni correction to control for multiple comparison errors (n=2, p<.025).

## 3. Results

### 3.1 Subjects data

#### 3.1.1 Demographic, self-report and brain volume data

The demographic and clinical data are shown in Table 1. Groups did not differ in terms of age as well as educational level. However, BMI of AN was significantly lower than in BN and CN, while BN and CN did not differ. Disease duration was similar for AN and BN. All EDI-2 scores were different among groups, except for EDI-2 Maturity Fear. Post-hoc analyses of the significant group effects showed differences in ED compared to CN for all the psychopathological scales with AN,BN>CN (AN=BN). The only exceptions were Body dissatisfaction and Ineffectiveness, with BN>AN>CN and Bulimia with BN>AN,CN (AN=CN). SCL-90 dimensions were all higher in ED compared to CN (see Supplementary Table S1).

**Table 1.**
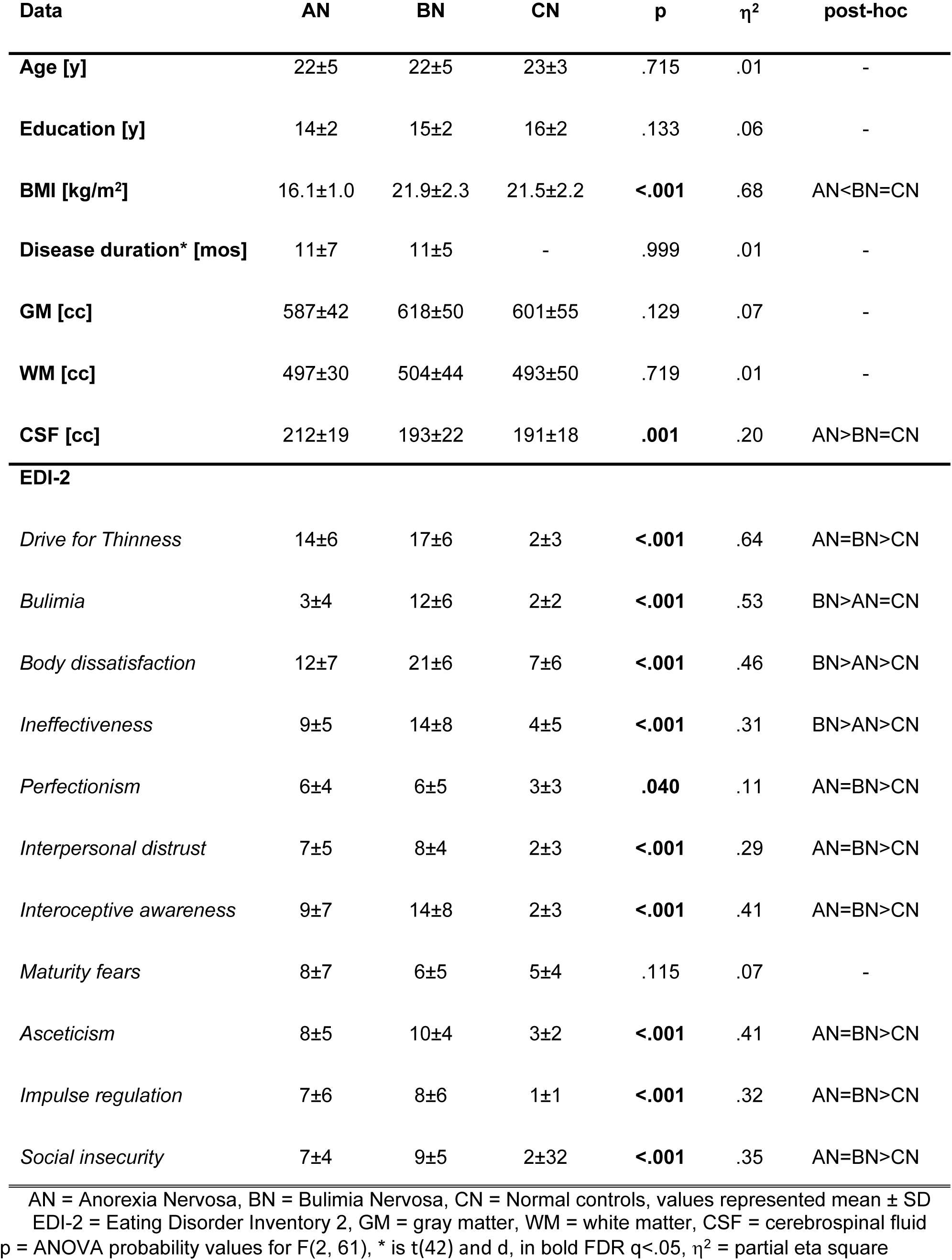
Demographic and clinical data separately for the three groups.

| Data | AN | BN | CN | p | $\eta^2$ | post-hoc |
| --- | --- | --- | --- | --- | --- | --- |
| Age [y] | 22±5 | 22±5 | 23±3 | .715 | .01 | - |
| Education [y] | 14±2 | 15±2 | 16±2 | .133 | .06 | - |
| BMI [kg/m<sup>2</sup>] | 16.1±1.0 | 21.9±2.3 | 21.5±2.2 | **<.001** | .68 | AN<BN=CN |
| **Disease duration\* [mos]** | 11±7 | 11±5 | - | .999 | .01 | - |
| **GM [cc]** | 587±42 | 618±50 | 601±55 | .129 | .07 | - |
| **WM [cc]** | 497±30 | 504±44 | 493±50 | .719 | .01 | - |
| **CSF [cc]** | 212±19 | 193±22 | 191±18 | **.001** | .20 | AN>BN=CN |
| **EDI-2** |  |  |  |  |  |  |
| *Drive for Thinness* | 14±6 | 17±6 | 2±3 | **<.001** | .64 | AN=BN>CN |
| *Bulimia* | 3±4 | 12±6 | 2±2 | **<.001** | .53 | BN>AN=CN |
| *Body dissatisfaction* | 12±7 | 21±6 | 7±6 | **<.001** | .46 | BN>AN>CN |
| *Ineffectiveness* | 9±5 | 14±8 | 4±5 | **<.001** | .31 | BN>AN>CN |
| *Perfectionism* | 6±4 | 6±5 | 3±3 | **.040** | .11 | AN=BN>CN |
| *Interpersonal distrust* | 7±5 | 8±4 | 2±3 | **<.001** | .29 | AN=BN>CN |
| *Interoceptive awareness* | 9±7 | 14±8 | 2±3 | **<.001** | .41 | AN=BN>CN |
| *Maturity fears* | 8±7 | 6±5 | 5±4 | .115 | .07 | - |
| *Asceticism* | 8±5 | 10±4 | 3±2 | **<.001** | .41 | AN=BN>CN |
| *Impulse regulation* | 7±6 | 8±6 | 1±1 | **<.001** | .32 | AN=BN>CN |
| *Social insecurity* | 7±4 | 9±5 | 2±3 | **<.001** | .35 | AN=BN>CN |

AN = Anorexia Nervosa, BN = Bulimia Nervosa, CN = Normal controls, values represented mean ± SD EDI-2 = Eating Disorder Inventory 2, GM = gray matter, WM = white matter, CSF = cerebrospinal fluid p = ANOVA probability values for F(2, 61), \* is t(42) and d, in bold FDR q<.05, $\eta^2$ = partial eta square

Self-reported empathy scores did not differ between groups (p=.153). The TAS-20 total score as well as the Difficulty in identifying and Difficulty in describing feelings subscales displayed increased alexithymia in AN and BN compared to CN (all p<.001), but no differences were evident between the two clinical groups (all p>.503).

For brain atrophy, the CSF global volume was increased in AN compared to both BN (p<.010) and CN (p<.002), while BN and CN did not differ (p<.895).

#### 3.1.2 Neuropsychological data

Regarding neuropsychological performance, a significant group effect emerged only for the Stroop Color-Word Test (p<.003). Post-hoc tests indicated that AN and BN showed greater interference than CN (AN>CN, p<.014; BN>CN, p<.004), while the clinical groups did not differ significantly (AN=BN, p<.812). For all other tasks, no significant group difference emerged (all FDR corrected p>.05). Please see Supplementary Table S2 for details.

#### 3.1.3 Behavioral performance during affective responsiveness task

The rmANOVA revealed a significant task effect (p<.001, η^2^=.46), but no main effect of group (p<.329) and no significant task-by-group interaction (p<.980). Post-hoc analyses of the significant emotion effect indicated that anger was the most difficult emotion to match (anger<fear, p<.001; anger<disgust, p<.001) and disgust the easiest (disgust>anger, p<.001; disgust>fear, p<.008). The control task was easier than all the emotion tasks (all p<0.001). Please see Table 2 for further details.

**Table 2.**
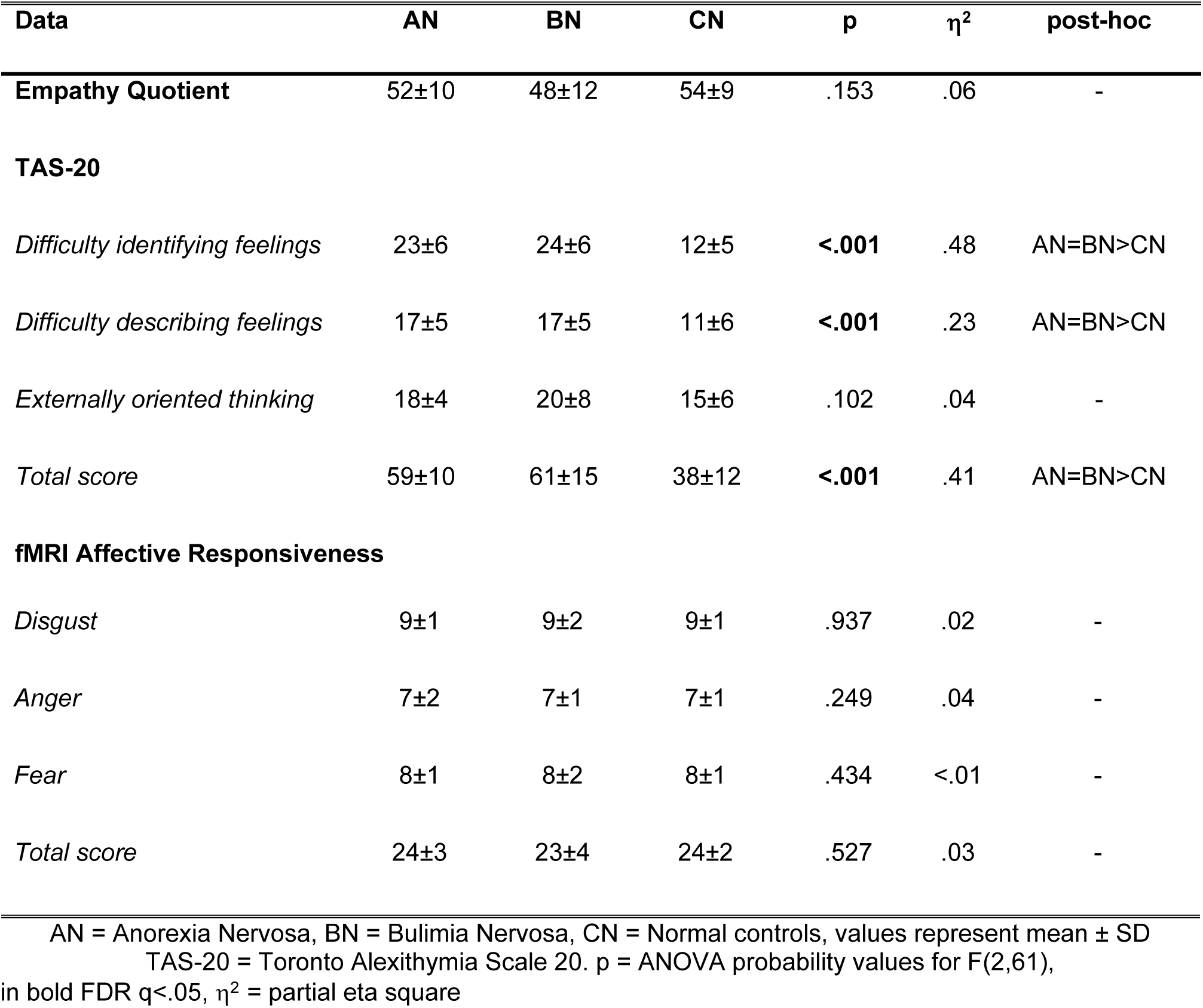
Self-reported empathy, alexithymia and behavioral performance during affective response task for the three groups.

| Data | AN | BN | CN | p | $\eta^2$ | post-hoc |
| --- | --- | --- | --- | --- | --- | --- |
| **Empathy Quotient** | 52±10 | 48±12 | 54±9 | .153 | .06 | - |
| **TAS-20** |  |  |  |  |  |  |
| *Difficulty identifying feelings* | 23±6 | 24±6 | 12±5 | **<.001** | .48 | AN=BN>CN |
| *Difficulty describing feelings* | 17±5 | 17±5 | 11±6 | **<.001** | .23 | AN=BN>CN |
| *Externally oriented thinking* | 18±4 | 20±8 | 15±6 | .102 | .04 | - |
| *Total score* | 59±10 | 61±15 | 38±12 | **<.001** | .41 | AN=BN>CN |
| fMRI Affective Responsiveness |  |  |  |  |  |  |
| *Disgust* | 9±1 | 9±2 | 9±1 | .937 | .02 | - |
| *Anger* | 7±2 | 7±1 | 7±1 | .249 | .04 | - |
| *Fear* | 8±1 | 8±2 | 8±1 | .434 | <.01 | - |
| *Total score* | 24±3 | 23±4 | 24±2 | .527 | .03 | - |

AN = Anorexia Nervosa, BN = Bulimia Nervosa, CN = Normal controls, values represent mean ± SD TAS-20 = Toronto Alexithymia Scale 20. p = ANOVA probability values for F(2,61), in bold FDR $q < .05$ , $\eta^2$ = partial eta square

### 3.2 Imaging negative emotional situations

#### 3.2.1 Whole-brain analyses

The contrasts for the group factor did not show significant activation differences in the brain.

#### 3.2.2 Amygdala ROI analysis

ROI analyses of the left and right amygdala revealed no significant differences between group, confirming whole-brain results (see Figure 2).

**Figure 2:**
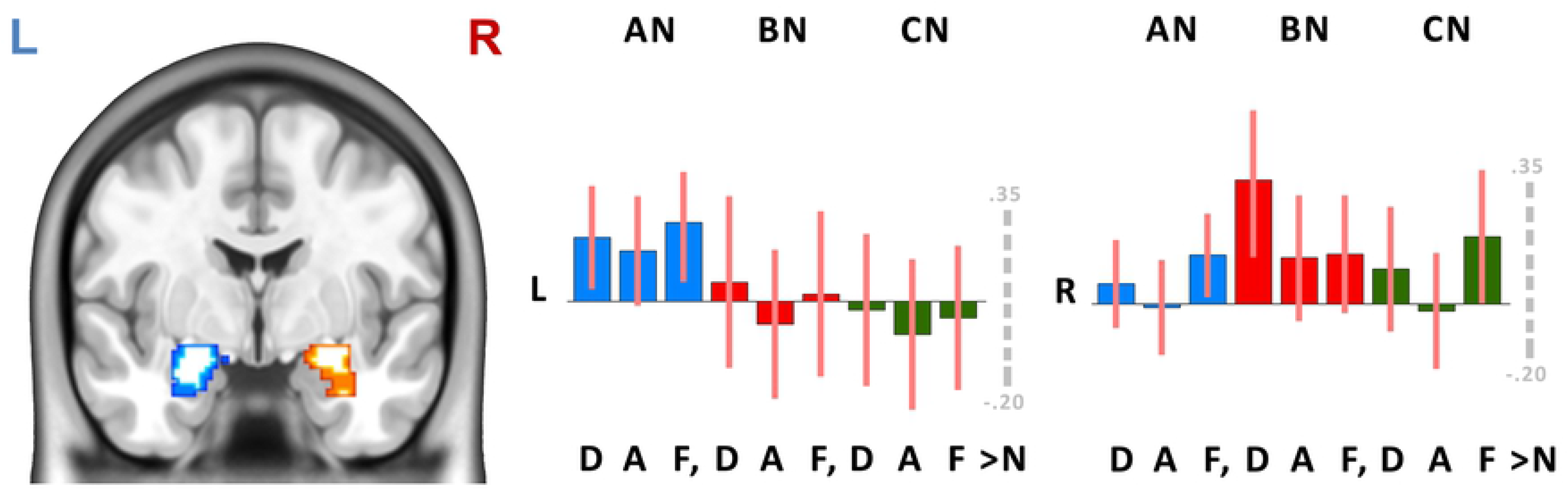
ROI analysis differences between groups. The figure plots the coefficients extracted from left and right amygdala during affective responsiveness fMRI for the different groups (AN = Anorexia Nervosa, BN = Bulimia Nervosa, CN = Normal controls, L = left, R = right, D = disgust, A = anger, F = fear, N = neutral). The stars indicate significant differences (p<.05), you can observe that no difference could be observed between groups in the left and right amygdala for negative emotions.

## 4. Discussion

This study’s main goal was to address affective responsiveness to negative emotional situations in treatment-naïve young females with AN or BN and matched controls. Thus, this study is one of the few that directly compare neuroimaging activation of different groups of ED with each other, thereby highlighting disorder-specific as well as transdiagnostic dysfunctional symptoms.

Based on previous literature we hypothesized to observe significant group differences in empathy and emotion processing between controls and ED. Instead, and in part unexpectedly, we detected mixed results: no significant group differences in behavioral performance or self-reported empathy, but significant differences in alexithymia. We also found no specific differences in neural activation for emotions. As expected, eating-specific and general psychopathology scales sharply distinguished participants affected by ED from healthy controls. Eating-specific scales like Body Dissatisfaction, Ineffectiveness and Bulimia further differentiated the clinical samples, with higher scores in BN.

### Neural network of affective responsiveness in ED

The emotional response paradigm we used activates consistently a set of brain regions [32–35], including the amygdala, which have been demonstrated to be involved in emotional empathy [23–25]. Interestingly, we did not observe a significant group effect for the amygdala or other brain areas. This lack of a significant group-by-emotion interaction in neural activation during our emotional responsiveness task is partly in line with previous reports of no group differences in neural activation between healthy controls and acute as well as recovered AN for explicit emotion recognition [17–22].

### Empathy and affective responsiveness in ED

Previous findings on empathy or emotion recognition are inconsistent: while some reported decreased self-reported emotional empathy but normal emotion recognition in acute AN [12], others observed no significant alteration in self-reported empathy in AN [13]. Additionally, a significant impairment in understanding others’ emotions and in the regulation of their own emotions has been reported in AN [40,41]. However, studies based on film clips did not find any specific abnormalities in emotion processing and emotional ratings, only attentional biases [42]. Some researchers pointed out that at least in a subgroup of individuals affected by BN, empathic mentalization was not impaired at all [43,44].

It has been hypothesized that the mixed results derived from the relatively poor sensitivity and specificity of the applied instruments, as already evidenced in the literature [30], but also considering our negative findings with a highly sensitive measurement of the brain activity it seems an incomplete explanation. A recent review [14] suggests an alternative explanation: ED individuals can recognize others’ basic emotions, but they lose this skill when emotions become more complex and are expressed within a relationship. Relevant to this kind of hypothesis, a dysfunctional middle prefrontal cortex (MPFC) activation has been reported for processing of visual stimuli depicting couples in intimate relationships in acute and recovered AN, pointing to a state-independent alteration of MPFC activation [45,46]. Additionally, the significantly elevated alexithymia scores in both clinical groups may be also associated with MPFC dysfunction, as this is an important hub for the interhemispheric integration and transfer [47]. Thus, MPFC function may be compromised in ED indicating a transdiagnostic alteration that seems especially relevant for the processing of complex stimuli, social functioning, and interpersonal interactions.

### 4.1 Limitations

Despite the novelty and relevance of this investigation of emotional responsiveness, the study had several limitations that need to be acknowledged. First, in principle, the fMRI task could be solved with different strategies, limiting the emotional response in favor of cognitive processing, but our finding of a similar activation of the amygdala for all groups limits the likelihood of this possibility. Second, we only investigated a small fragment of emotion processing and social cognition. More distinct differences between ED and controls could emerge if other processes that vary in complexity would be studied like affective vs. cognitive empathy or interpersonal affectivity and reactivity. Finally, different subtypes of participants (purging, non-purging) were included, but subsamples were too small to analyze their specific contributions.

## 5. Conclusions

Relying on self-report data and an affective responsiveness task, we must argue against the hypothesis that participants with ED display reduced emotional responsiveness. This supports the hypothesis that relational difficulties, as well as therapeutic resistance, are not secondary to simple difficulty in feeling and identifying basic negative emotions in first episode, treatment-naive AN and BN participants.

According to the theories which link the pathogenesis of anorexia nervosa to attachment processes [48,49], the present findings support the hypothesis that the difficulty in building meaningful relationships which characterize young women with ED may be more related to attachment issues based on early attachment experiences rather than from current difficulties in the emotional responsiveness to their own and others emotions.

## Acknowledgements

The authors declare no conflicts of interest. This research received no specific grant from any funding agency, commercial or not-for-profit sectors.

The data that support the findings of this study are available from the authors, without undue reservation, to any qualified researcher.

## Authors contributions

Conception and design of the study: FA, FDA, PC, LL, GAD, BD, SF.

Acquisition and analysis of data: FDA, AS, PC, LL, MB, AC.

Drafting the manuscript or figures: FA, FDA, AS, PC, LL, GAD, AB, MB, AC, BV, BD, SF.

